# Scaling accurate genetic variant discovery to tens of thousands of samples

**DOI:** 10.1101/201178

**Authors:** Ryan Poplin, Valentin Ruano-Rubio, Mark A. DePristo, Tim J. Fennell, Mauricio O. Carneiro, Geraldine A. Van der Auwera, David E. Kling, Laura D. Gauthier, Ami Levy-Moonshine, David Roazen, Khalid Shakir, Joel Thibault, Sheila Chandran, Chris Whelan, Monkol Lek, Stacey Gabriel, Mark J Daly, Ben Neale, Daniel G. MacArthur, Eric Banks

## Abstract

Comprehensive disease gene discovery in both common and rare diseases will require the efficient and accurate detection of all classes of genetic variation across tens to hundreds of thousands of human samples. We describe here a novel assembly-based approach to variant calling, the GATK HaplotypeCaller (HC) and Reference Confidence Model (RCM), that determines genotype likelihoods independently per-sample but performs joint calling across all samples within a project simultaneously. We show by calling over 90,000 samples from the Exome Aggregation Consortium (ExAC) that, in contrast to other algorithms, the HC-RCM scales efficiently to very large sample sizes without loss in accuracy; and that the accuracy of indel variant calling is superior in comparison to other algorithms. More importantly, the HC-RCM produces a fully squared-off matrix of genotypes across all samples at every genomic position being investigated. The HC-RCM is a novel, scalable, assembly-based algorithm with abundant applications for population genetics and clinical studies.

## 1 Introduction

Next-generation sequencing technologies combined with large population cohorts are facilitating the identification of rare allelic variants for population and medical genetic studies [18, 4, 25, 14]. Although high-throughput sequencing platforms currently generate up to four billion paired-end reads per run, low signal-to-noise ratios necessitate the use of data processing algorithms that differentiate true variants from machine-generated artifacts [19, 11].

The first generation of variant calling algorithms scanned short read data mapped to a reference sequence to identify mismatches [1, 6, 15, 17, 22]. Reference sequence mismatches were probabilistically sorted using algorithms that took into account the reported base quality and context prior to calling variants [22]. These position-based or so-called “pileup” callers, which include SAMtools Li et al 2009 and the GATK’s UnifiedGenotyper (UG), proved highly effective at calling small nucleotide polymorphisms (SNPs) but are unable to attain high accuracy for indel variants due to their reliance on independent short read alignments to a reference sequence [1, 6, 15, 17, 22]. In contrast, assembly-based variant callers including Platypus [22] and the GATK’s HaplotypeCaller (HC), construct theoretical haplotypes via de Bruijn-like graphs from a consensus of the reads covering the genomic region [13, 22, 7, 10, 20]. Although assembly-based algorithms call indels with greater accuracy, they do not scale well due in part to exponential increases in graph complexity with the number of samples. Thus, calling variants jointly on large numbers of samples becomes increasingly computationally intensive until the requirements exceed hardware performance limitations [6].

One naive solution would be simply to discover and genotype variants in each sample separately and then merge the independently discovered variants across all samples. In this way, data from a single sample would only be retained at positions in which that sample is variant. A major shortcoming of this approach is the only genotype calls present would be heterozygous or homozygous variant. It would be impossible to determine whether missing genotype calls were homozygous reference or void of any read data at all. As an illustration of this shortcoming, take the case where a mutation is discovered in a single sample within a cohort; a naively merged list would merely contain “no-call” genotypes for all other samples making estimating the population allele frequency difficult. What is required for the most accurate estimates of population allele frequencies is instead to jointly call variants over all samples together, creating a “squared-off matrix” of samples by genomic position where each cell in the matrix contains genotype likelihoods for all possible alleles (including the reference) at the corresponding genomic position. In addition, is has been previously shown [6] that joint calling has several other advantages: it improves sensitivity at low coverage positions and powers the training of a more accurate statistical filtering model, which is a crucial step in the variant discovery pipeline (Figure 1).

**Figure 1:**
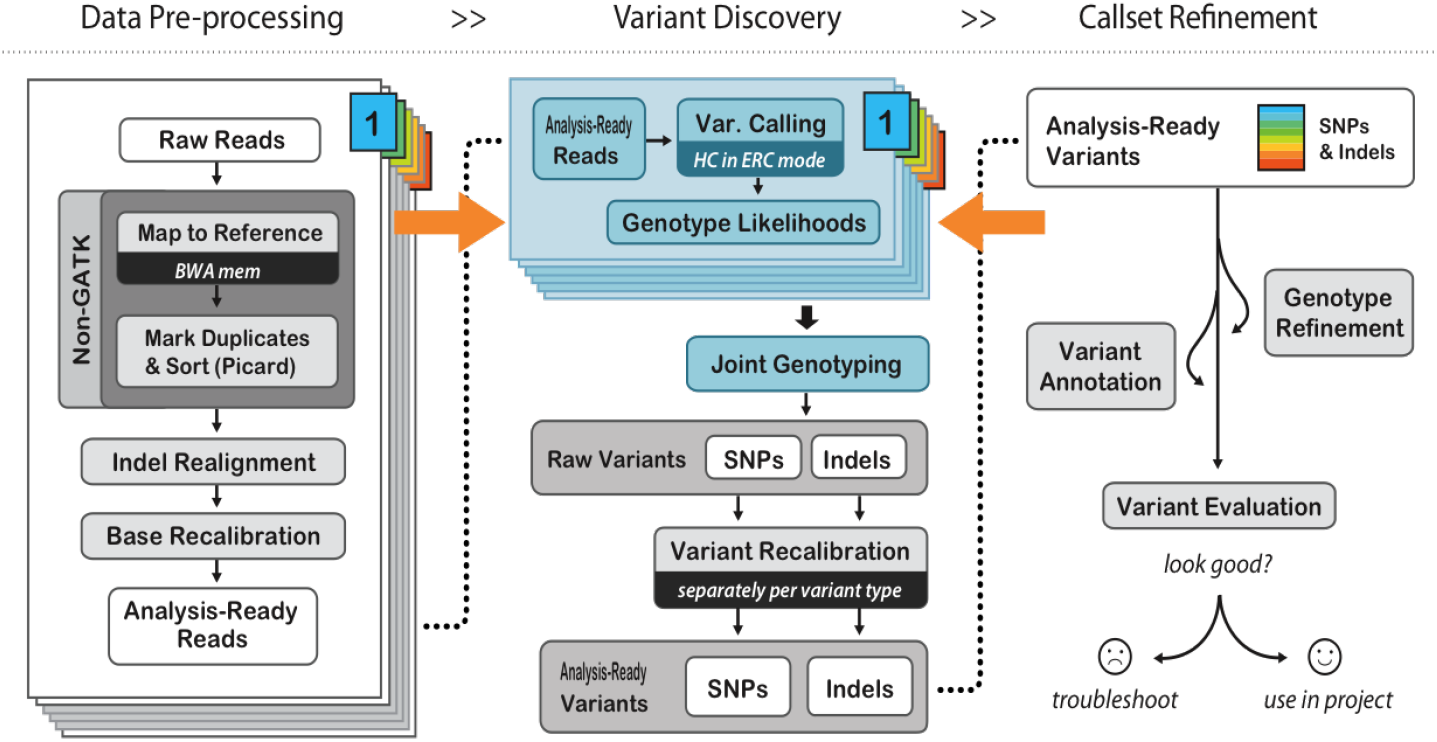
GATK Variant Calling Best Practices. The HaplotypeCaller takes in analysis-ready reads and performs variant calling per sample to produce unfiltered genotype likelihoods.

To solve the problem of joint calling large cohorts using graph-based assembly without introducing intractable computational complexity, the Reference Confidence Model was developed as a module for the HC (Figure 2). In brief, this combined HaplotypeCaller-Reference Confidence Model (HC-RCM) algorithm constructs candidate haplotypes for each individual sample in the cohort, avoiding the exponential complexity of a graph describing all samples simultaneously and ultimately accelerating variant calling compared with the previous approach to joint calling. Constructed haplotypes are used to calculate genotype likelihoods using a pair-hidden Markov model (pair-HMM), and these likelihoods are stored in an intermediate file for subsequent joint variant calling across all samples (which includes allele frequency estimation and genotype assignment). In addition to likelihoods for all alleles explicitly observed a sample’s reads, the model generates the likelihood over the set of unobserved, non-reference alleles. Here it is shown that the accuracy of the HC-RCM algorithm is comparable to or better than other widely used tools and that the computational performance on large sample sizes is superior.

**Figure 2:**
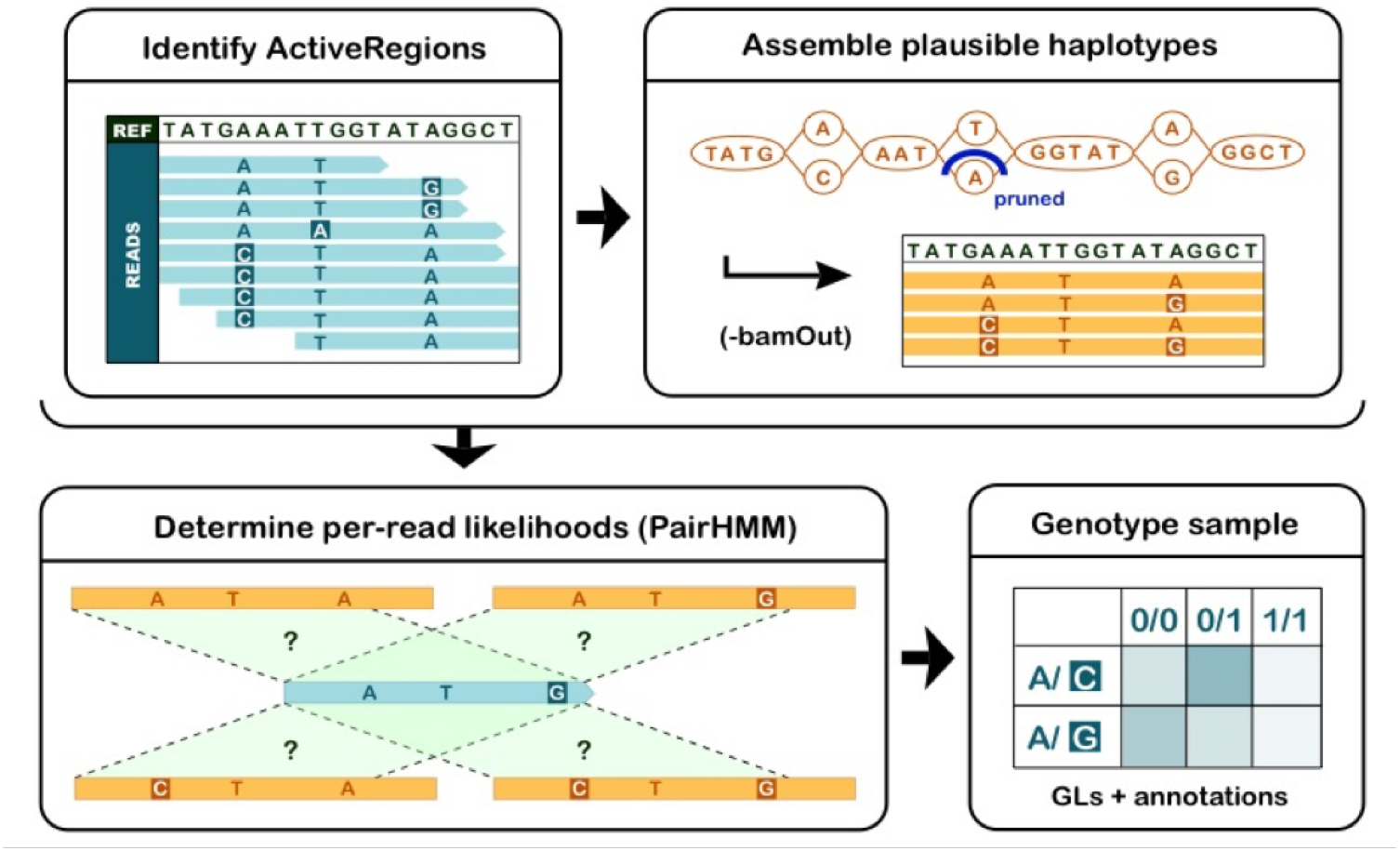
HaplotypeCaller-Reference Confidence Model overview. The four basic steps of variant calling with the HC-RCM including identification of the ActiveRegions, assembly of candidate haplotypes using de Bruijn-like graphs, determination of the per-read likelihoods of candidate haplotypes using a pair-HMM, and genotype assignment.

## 2 Results

### 2.1 Brief Methodology

The role of the HaplotypeCaller within the GATK variant calling pipeline is shown in Figure 1 (see http://www.broadinstitute.org/gatk for more details regarding the GATK Best Practices). High-throughput sequencing analyses start with raw read data (FASTQ files) that are converted to recalibrated, analysis-ready reads in the Binary sequence Alignment/Map (BAM) format using SAMtools and GATK modules [16, 6]. Reads are initially aligned to a reference sequence e.g. human reference genome (GRCh37) using an mapping/aligner program e.g. Burrows-Wheeler Aligner (BWA) [16, 5].

For computational efficiency, variant calling is focused on “ActiveRegions”, or regions of the genome that vary significantly from the reference. ActiveRegions are defined by genomic intervals where the BWA-aligned reads contain evidence that they are in disagreement with the reference, using criteria such as base mismatches, insertion/deletion markers in the reads and high base-quality soft clips [16]. The reads from these ActiveRegions are split into overlapping subsequences, or k-mers, and subsequently reassembled using de-Bruijn-like graphs into candidate haplotypes. This is followed by the construction of a pair-HMM using state transition probabilities derived from the read base qualities. This pair-HMM is then used to calculate the likelihood that each read was derived from each haplotype (Supplemental Figure 5). These likelihoods are used to calculate the raw genotype likelihoods for each candidate variant. Ultimately, genotype likelihoods across all samples are used to call raw variants for the cohort.

The raw, unfiltered output of the HaplotypeCaller is not appropriate to be used for downstream analyses. The HC-RCM aims to attain maximum sensitivity with the consequence of retaining some false positive variants. The subsequent steps of the GATK Best Practices pipeline (Figure 1) identify and filter out false positives to achieve maximum accuracy, as described in previous in previous work [6]. For the accuracy results presented here, the HC-RCM output is subjected to joint genotyping and variant recalibration in accordance with the GATK Best Practices.

For further details on the HC-RCM algorithm, see the Supplemental Methods section.

### 2.2 Comparative Analyses

To validate the sensitivity and specificity of this new variant calling algorithm, a set of analyses was performed comparing the HaplotypeCaller with three other variant calling algorithms (Figure 3). These other algorithms consisted of two pileup variant callers, SAMtools[17] and the GATK Unified Genotyper (UG) [6], as well as Platypus, another assembly-based algorithm [22]. SNPs and indels were called with each of the four variant calling algorithms from whole genome and exome-capture sequencing data from the well characterized CEU HapMap trio [2]. Variant caller accuracy was evaluated using the Genome in a Bottle standard (GiaB) [26] as described in the supplemental methods.

**Figure 3:**
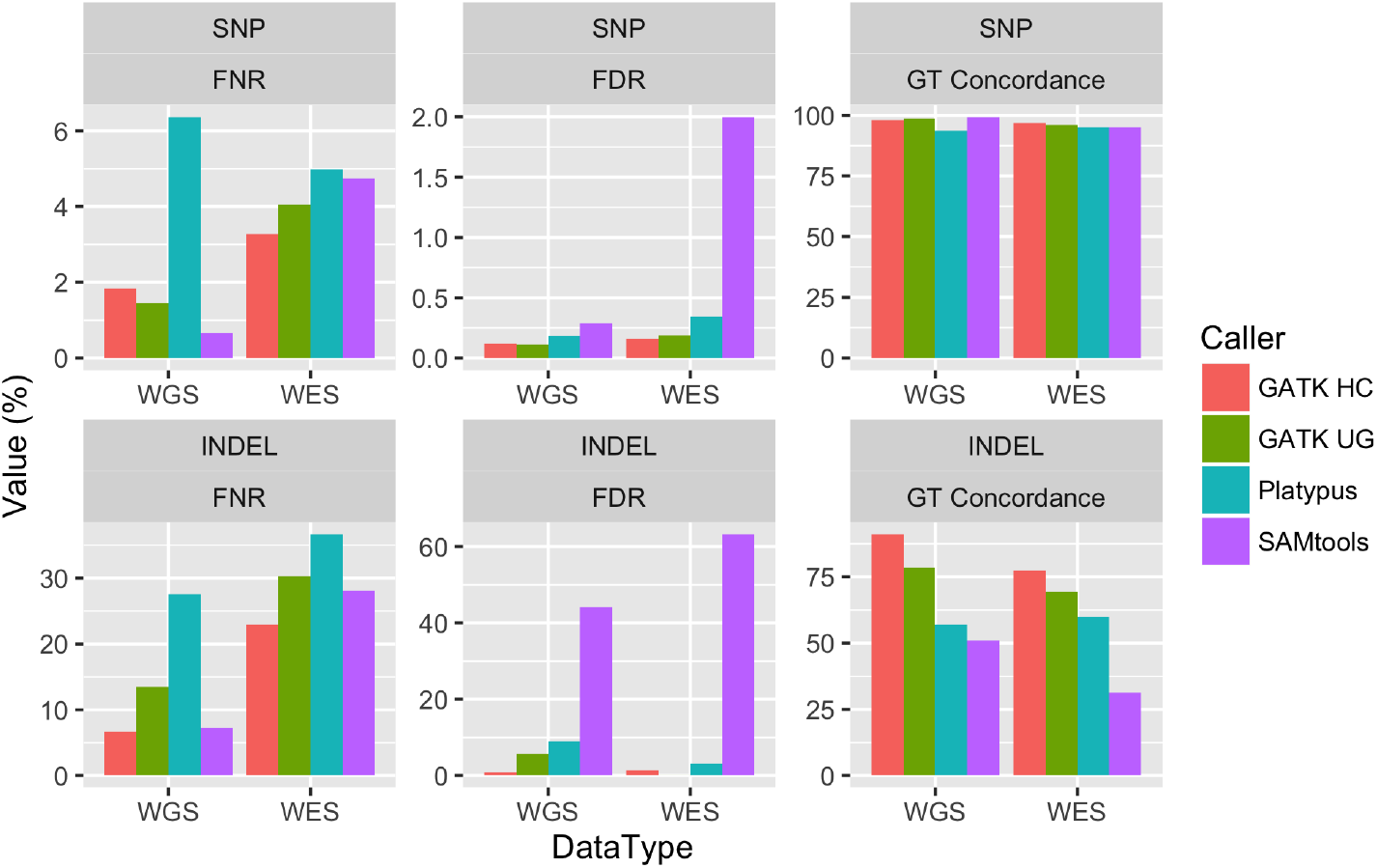
Variant caller comparison. Accuracy comparisons of both SNP and indel variant calls over all three samples in the CEU trio by the GATK HaplotypeCaller, Platypus, SAMtools, and Unified Genotyper. WGS data was PCR-free, 250bp paired-end reads sequenced by an Illumina HiSeq. WES data was 76bp paired-end reads sequenced by an Illumina HiSeq. Sensitivity is plotted as false negative rate (FNR), for which lower values equate to better sensitivity. Specificity is plotted as false discovery rate FDR, for which lower values are also better. For genotype concordance higher values indicate better genotype call accuracy.

Figure 3 shows the results of these analyses, which indicate that the HaplotypeCaller effectively calls both SNP and indel variants from both whole genome and exome-captured sequence data. Although Figure 3 shows that SAMtools called a greater number of both SNP and indel variants compared to the other callers, the specificity of these calls was substantially lower than the other algorithms, especially for indel variants, as indicated by high false discovery rates (FDR). In contrast, the HaplotypeCaller algorithm called large numbers of SNP and indel variants with high sensitivity and low FDRs on both whole genome and exome captured DNA, suggesting that the HaplotypeCaller has both high sensitivity and specificity.

To determine algorithm genotype assignment accuracy, the genotype concordance was determined. Genotype concordance measures the accuracy of genotype assignments of true positive variants to a gold standard (GiaB). In contrast to the other variant calling algorithms, the HaplotypeCaller had exceptionally high genotype concordance values for indel variant calls from both whole genome and exome captured data. These results suggest that the HaplotypeCaller calls genotypes with superior accuracy compared with the other algorithms.

### 2.3 Scaling

The Reference Confidence Model was integrated into the HaplotypeCaller to enable scaling of joint calling up to hundreds of thousands of exomes. To determine how this algorithm performs on large sample sets, runtime was examined as a function of the number of samples. Figure 4 shows the runtimes for variant calling using GATK HaplotypeCaller, GATK UnifiedGenotyper, Platypus and SAMtools. For the 20X exomes used here, HaplotypeCaller requires slightly more CPU-hours than the other algorithms for up to 250 samples. Beyond 250 samples, HaplotypeCaller runtime continues to increase linearly with the number of samples, while the other algorithms increase superlinearly. Furthermore, the efficient scaling of the HC-RCM algorithm was leveraged to produce a joint call set with over 90,000 exomes for the Exome Aggregation Consortium [14].

**Figure 4:**
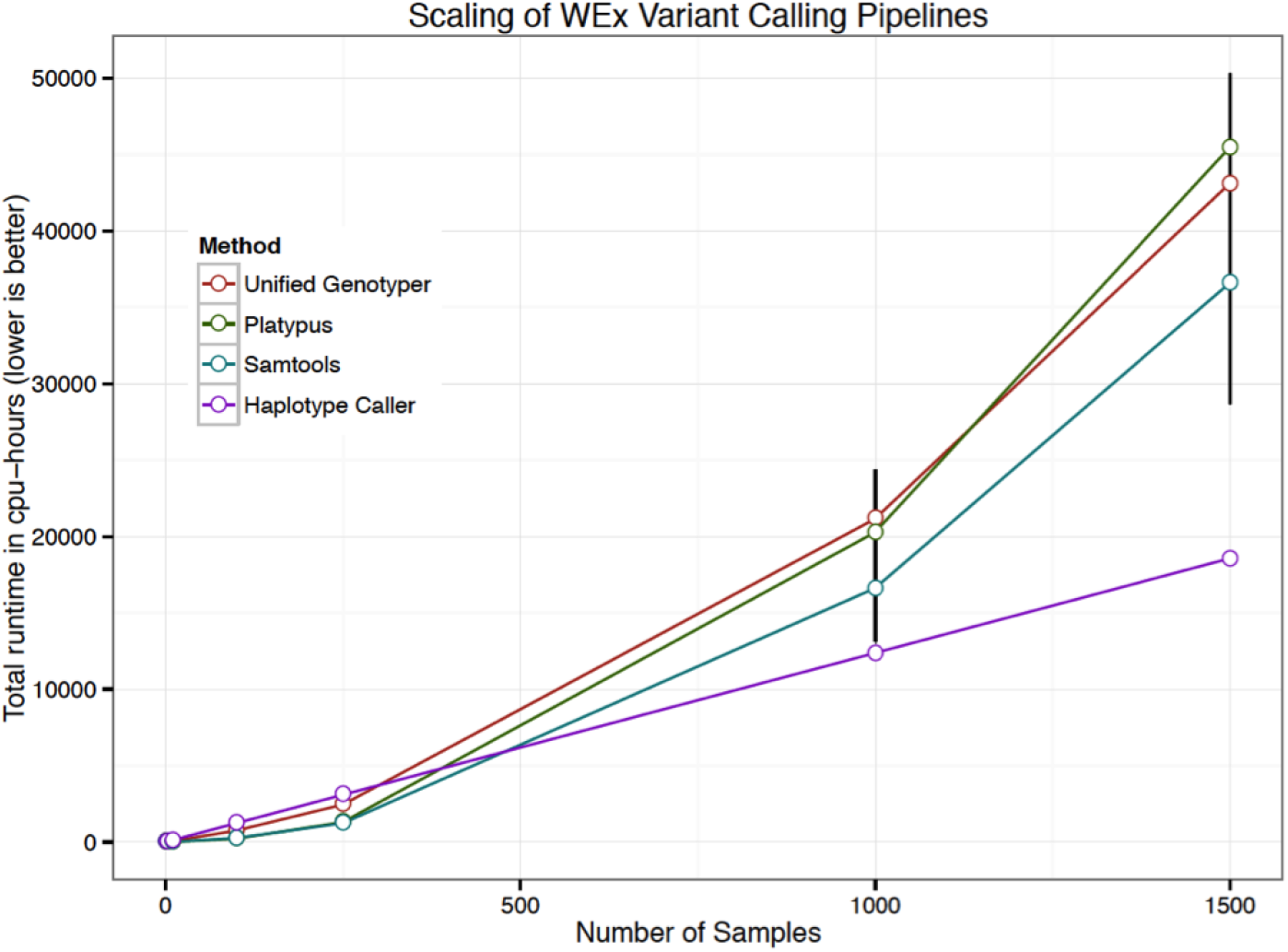
Scaling of whole exome calling with the HaplotypeCaller-Reference Confidence Model. Comparison of computational runtimes as a function of sample size. For each algorithm, variants were called over 1M bases of genomic territory and extrapolated to full exome runtime. Error bars represent 95% confidence intervals from 10 independent runs.

## 3 Discussion

The above validation studies demonstrate that the HaplotypeCaller calls both SNP and indel variants with high sensitivity and specificity. Now part of the widely used GATK pipeline, the HC removes a critical bottleneck in variant calling by enabling this algorithm to scale without losses in accuracy or sensitivity.

Our analyses show that SAMtools is the most sensitive SNP variant caller using both whole genome and exome-captured data. These results are consistent with other studies for whole genomic data which have shown that locus-based algorithms accurately call SNP variants [17, 22]. However, SAMtools also has a high false discovery rate for SNP variant calls, suggesting that though highly sensitive, it lacks the specificity of the other variant calling algorithms evaluated in this study.

In our results, the HaplotypeCaller was the most sensitive indel variant caller using both whole genome and exome-capture data, which is a conclusion consistent with other reports comparing variant callers [22, 21, 9]. However, in contrast with the report of Rimmer, et al., the results here show that the HaplotypeCaller had the lowest FDR on whole genomic data, suggesting that it had the highest specificity for indel variant calls [22]. Reasons for the discrepancies between the two studies, include different versions of the algorithm used; the current study used the HaplotypeCaller (v3.4) while Rimmer, et al used a much older version (v2.5). It is likely that more recent versions of the HaplotypeCaller software improved variant calling accuracy as has been reported by Narzisi, et al. [20] and our internal tests (data not shown). In addition to the different software versions used, there is a substantial difference in the validation methods between the two studies. Although Rimmer et al., used a fosmid dataset to validate their variant calls, the majority of variants called as false-positives by Platypus were actually present in dbSNP137, suggesting that their “truth” data contained a high frequency of errors [22]. In contrast, this analysis used the GiaB standard which contains variants validated using multiple data sets, platforms, variant callers, and filtering strategies suggesting that the GiaB is a highly validated “truth” dataset [26]. Using this high quality “truth” set it is demonstrated that the HaplotypeCaller calls indel variants with both high sensitivity and specificity.

These comparison data show that all of the algorithms assign genotypes with similar accuracy from SNP variants, consistent with the report of [22]. However, substantial differences in genotype concordance were observed among algorithms for indel variants. HaplotypeCaller’s local assembly and pairHMM likelihood calculation are more robust to errors in repetitive contexts, leading to the higher genotyping accuracy.

Platypus FDRs were higher than expected for both SNPs and indels in the WGS results. The orthogonal Illumina Platinum Genomes evaluation [9] showed a FDR of less than 1 While variant calling accuracy on a single sample is of primary importance in some studies, in nearly all contexts the interpretation of variants requires data derived from many samples. Analyses like genome-wide association studies leverage larger numbers of samples to increase their power to detect genotype-phenotype associations. In a clinical research setting, analysts often interpret the effect of a given putatively causal mutation based on population frequency estimated from a large joint call set of samples from healthy individuals used as a reference panel. To produce such a reference panel, it is crucial that all samples be jointly called and evaluated across as many other samples as possible. The estimation of the frequencies of rare alleles in large populations is possible only with accurate and confident genotype calls for all samples, especially those that are homozygous for the reference allele.

Joint calling increases variant calling sensitivity over low coverage regions and improves filtering accuracy [6]. The HaplotypeCaller algorithm further improves accuracy by using a local assembly method for variant discovery. However joint calling across large sets of samples requires an algorithm that can effectively scale. HC-RCM runtimes increase linearly with sample number, enabling the algorithm to produce data across tens of thousands of samples, as demonstrated by the production of a call set featuring over 90,000 exome samples used in the ExAC study [14].

## 4 Supplemental Methods

### 4.1 Data

Input data for the variant caller evaluations were obtained from the 1000 genomes repository [5]. Both data sets, consisting of the 2×250bp PCR-free whole genome samples and 2×76bp exome samples, were retrieved as BAM files of high coverage CEU trio samples. These samples, NA12891, NA12892, NA12878 representing the father, mother, and daughter respectively have been described previously [2]. Prior to analysis, all sequencing data were aligned with the Burrows-Wheeler transform algorithm (BWA) [16]. The Genome in a Bottle (GiaB) standard version 2.18 for NA12878 was used for the variant caller accuracy comparison [26].

### 4.2 Software

Software versions were Platypus version 0.7.8, SAMtools version 1.1 and the UnifiedGenotyper and HaplotypeCaller-Reference Confidence Model (HC-RCM) were obtained from the Genome Analysis Toolkit version 3.4. The authors made a good faith effort to run each tool according to its respective best practice recommendations by following the directions found on each tool author’s webpage.

### 4.3 Evaluation Metrics

Figure 3 reports variant caller accuracy using false negative rate (FNR), false discovery rate (FDR), and genotype concordance with respect to the Genome in a Bottle (GiaB) reference standard as the truth set [26]. A variant at a position listed in the truth set that also has an alternate allele matching the truth set is considered a true positive (TP). A variant that is called within the GiaB confidence region that does not occur in the list of truth variants or have an alternate allele matching the truth variant is considered a false positive (FP).

Algorithm sensitivity is calculated by dividing the number of true positive variants by the total number of variants in the GiaB standard [26, 22]. Here we report the FNR (1-sensitivity) to better visualize the differences between algorithms.

The FDR describes the frequency of incorrect variant calls in a call set and is calculated by dividing the number of false-positive (FP) variant calls by total number of called variants (the sum of both true and false positive variant calls) as described [22, 6]. Genotype concordance (GT concordance), measures the accuracy of genotype calls and is calculated by dividing the number of correctly assigned genotypes (both heterozygous and homozygous variant) by the total number of true positive variants [26].

### 4.4 HaplotypeCaller Algorithm

HaplotypeCaller is the method responsible for variant calling in the current best practices version of the GATK pipeline for the processing of high-throughput sequencing data. This component was developed to address the shortcomings of previous locus-based callers, primarily an inability to call structural variants including indels and repetitive sequences [22]. The HaplotypeCaller’s approach to variant calling is reference-based local reassembly of genomic regions containing non-reference evidence. Using this approach, the method presented here avoids many of the pitfalls of global alignment to the reference sequence used in mapping/locus-based approaches e.g. SAMtools and the Unified Genotyper [17, 6]. Given that this local assembly approach becomes exponentially more computationally intensive with increasing numbers of samples, the reference confidence model (RCM) was developed to reduce the computational burden associated with the tandem local reassembly of multiple samples. The combined algorithm is described in detail below. There are several preprocessing steps (Figure 1) that are explained in detail on the GATK website www.broadinstitute.org/gatk and in previous publications [6, 3]. Initially, raw sequence data is aligned and de-duplicated using BWA and Picard Tools, respectively [17, 15]. Additional preprocessing steps used to produce analysis-ready reads, including the Base Quality Score Recalibrator (BQSR), are part of the GATK pipeline and have been described previously [6, 3]. Note also that reads with mapping quality less than 20 are filtered by the tool engine and excluded from HaplotypeCaller.

#### 4.4.1 Defining ActiveRegions

To define an ActiveRegion of a sample’s genome using data from that sample’s reads, the HaplotypeCaller operates in three phases. It computes an “active probability” for each locus, smoothes the probability signal via convolution, and thresholds the resulting signal to define the ActiveRegion boundaries. Using the original alignment assigned by BWA, the HaplotypeCaller performs genotyping at each pileup position comparing the reference allele with any non-reference possibility, incorporating mismatch evidence, but also insertions, deletions, and soft-clips. Sites are assigned an “active probability” based on these genotype likelihoods and a heterozygosity prior (0.001 for SNPs and 0.0001 for indels by default). That probability is then convolved across loci with a Gaussian kernel (sigma of 17bp by default) to expand the activity signal. An ActiveRegion is thus defined as an interval of contiguous loci where the active probability exceeds a certain threshold (0.002 by default.) By default, HaplotypeCaller incorporates data from reads that cover an interval of up to 100bp adjacent to but outside the thresholded ActiveRegion during assembly, but these reads do not contribute to genotype likelihoods. Reads outside the ActiveRegion and its extension are not considered at all. Minimum ActiveRegion size is by default 50bp while the maximum size is 300bp. If an ActiveRegion exceeds the maximum size after the thresholding step then is it split into two, such that the regions for which each will emit variants abut, but reads overlapping the junction will be considered in both.

#### 4.4.2 Graph Construction

The second step involves de Bruijn-like graph construction for each ActiveRegion [23]. The reads within the current ActiveRegion and the reference sequence corresponding to that ActiveRegion are parsed into k-mers of 10 and 25 nucleotides in length. If the k-mer-ized reference sequence contains nonunique k-mers, then the value k in incremented by 10 until a maximum of 65. If the reference sequence for the ActiveRegion contains non-unique 65-mers, then assembly is aborted. The graph is initialized by connecting overlapping reference k-mers into a single path. Edges in the graph are assigned weights according to how many reads contain each pair of k-mers the edge connects. The graph is simplified by merging paths with entire k-mers in common. Edges with limited k-mer support are pruned out of the graph and potential haplotypes are removed if supported by fewer than two reads by default. Paths that do not have a terminal kmer that connects back to the reference haplotype, referred to as “dangling tails”, are attempted to be merged to the reference using Smith-Waterman alignment [24]. If no overlap with the reference is found, paths containing the dangling tail will be discarded. Each candidate haplotype is aligned to the reference sequence using Smith-Waterman. The output of this alignment generates a CIGAR string for each candidate haplotype (*H_j_*) [17]. This CIGAR is used to help translate the haplotypes into the variants that will be output in the VCF.

#### 4.4.3 Pair-HMM

A paired Hidden Markov Model (pair-HMM) is used to determine the likelihood for each combination of candidate haplotype (*H_j_*) and read data (*R_i_*), namely *P* (*R_i_*|*H_j_*) [12, 8].

Figure 5 shows a graphical representation of a global alignment model, which is a simplified version of the pair-HMM used herein. For additional details and a description of the complete model, the reader is referred to Durbin et al., [8]. The pair-HMM calculates an alignment score for each read (*R_i_*) and candidate haplotype (*H_j_*). There are three primary “states” of the algorithm, match (*M_ij_*), insertion (*I_ij_*), and deletion (*D_ij_*). An alignment of a read to a haplotype can be described by a sequence of these states. The transition probability from the match to the insertion or deletion state is given by the gap open penalty (*δ*). The probability that an alignment stays in either the insertion or deletion state is given by the gap extension penalty (*ϵ*). The gap extension penalty is held constant (default 10) while the gap open penalty may be derived from GATK’s BQSR if base insertion and deletion qualities have been output in the recalibrated BAM. If BQSR base insertion and deletion qualities are not available, a constant value of 45 is assigned. For alignments in the match state, the probability of emitting a base identical to the reference (*P_ij_*) is given by the complement of the base error probability given by the base quality.

**Figure 5:**
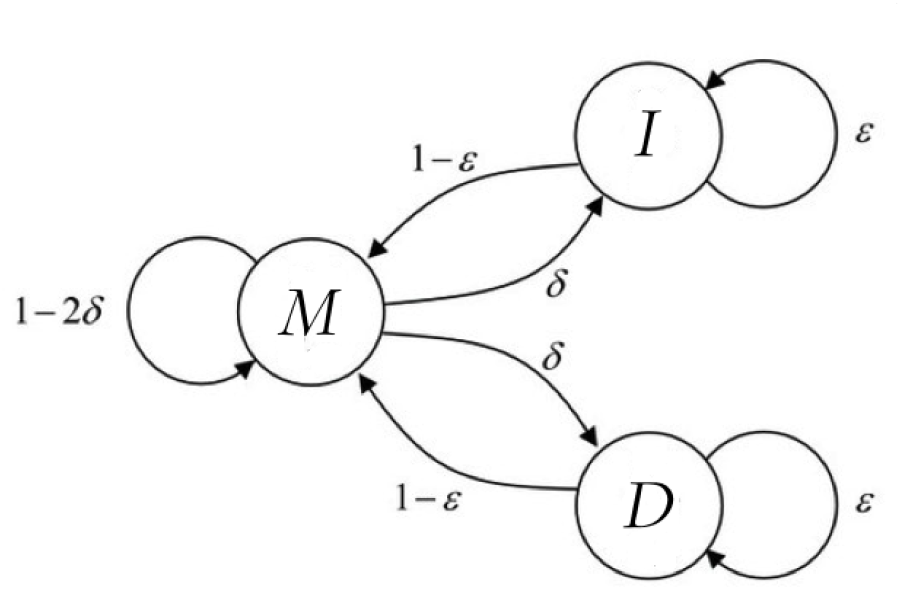
Pair-HMM overview. This diagram shown depicts the states, transitions, and transition rates for the read-haplotype alignment. The alignment of a read to a candidate haplotype is allowed to take on one of three states; match, insertion, or deletion (M, I, or D). *ϵ* represents the gap extension penalty while *δ* is the gap open penalty. Other transition probabilities are defined based on the constraint that the sum of the transition probabilities must equal one.

Recurrence relations for the states with respect to position i in the read and position j in the haplotype are given as follows:

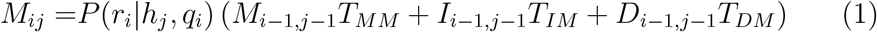

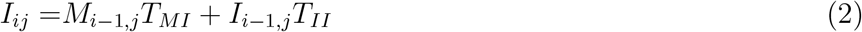

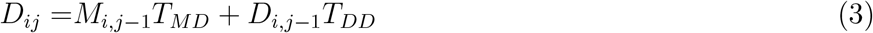

where *T_MI_* = *T_MD_* = *δ T_MM_* = 1 – 2*δ, T_II_* = *T_DD_* = *ϵ, T_IM_* = *T_DM_* = 1 – *ϵ* and *T_ID_* = *T_DI_* = 0

The pair-HMM can be used to correct for PCR errors that can lead to spurious indel calls. In this error correction mode, when the alignment sequence is being calculated for a tandem repeat sequence in the reference, the base insertion and deletion qualities are decreased in order to convey the reduced confidence in indels occurring within repetitive contexts. The default is to use a conservative correction. For PCR-free genomes, the authors recommend setting the PCR indel model to “NONE”.

#### 4.4.4 Assigning Genotypes

The CIGAR derived from the Smith-Waterman alignment of the discovered haplotypes is used to translate the haplotypes into variant events with respect to the reference. Genotyping is then performed at each of the events discovered in any haplotype. Per-read haplotype likelihood output from the pair-HMM is used to calculate raw genotype likelihoods using a Bayesian model. Given that multiple haplotypes may support a variant allele, the per-read allele likelihood at a given variant position is taken as the maximum of the likelihoods for haplotypes containing the allele. Equation 1 gives the genotype likelihood for a diploid genotype *G_i_* composed of one copy of allele *A*_1_ and one copy of allele *A*_2_. Briefly, the probability of a candidate genotype *P*(*R_i_|G_l_*), is the product over all the reads of the mean read likelihoods for the alleles in the specified genotype

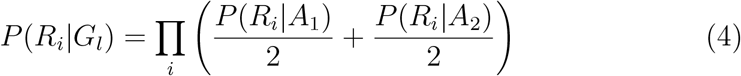

To determine the posterior probability of each candidate genotype *P*(*G_l_|R_i_*), the raw genotype likelihoods *P*(*R_i_|G_l_*) are marginalized using a Bayesian model:.

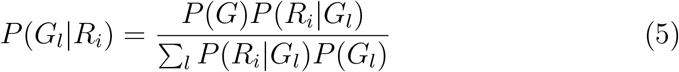

The numerator consists of the product of the prior probability of a genotype *P*(*G*) and a raw genotype likelihood divided by the sum of the likelihoods of all the possible genotypes for the set of alleles called in the variant. At this stage in the best practices pipeline, the genotype prior is flat such that the prior probability of all genotypes are equal. The GATK tool CalculateGenotypePosteriors can be used in post-processing to apply more informed priors.

#### 4.4.5 Reference Confidence Model

The two main requirements for joint calling large cohorts using local assembly are maintaining manageable compute complexity and outputting data for every variant in every sample. The reference confidence model addresses the former requirement by processing the samples individually, reducing the number of probable paths through the assembly graph, resulting in fewer candidate haplotypes, and decreasing the number of computationally expensive likelihood calculations that must be performed for each sample. The solution to the latter involves the addition of the symbolic non-reference allele `(<NON_REF>)` to aggregate evidence for a variant that was not explicitly called as an alternate allele. This additional candidate allele category enables the estimation of likelihoods to genotypes featuring an allele not seen in the sample in question. For variant sites, for each read the non-reference likelihood is set to the median of the set of allele likelihoods that are worst than the best allele. For sites that appear to be reference, non-reference likelihoods are estimated using a SNP model and an indel model.

Under the SNP model, base qualities are used to assign oer-read allele likelihoods for the reference and non-reference alleles. The per-read allele likelihoods for reference bases are assigned using the base error probability (*ϵ*) derived from the base quality. Bases that do not match the reference or are adjacent to soft clips, insertions, or deletions are considered non-reference evidence. The likelihood for read *R_i_* with base *b_i_* and base quality *q* compared to the reference allele with base *b_r_* is given by:

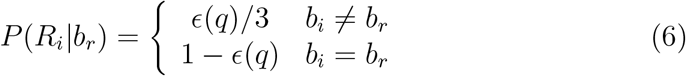

Where *ϵ* = 10^−*q*/10^ for base quality q. Deletions are assigned a constant quality of 30. Bases with base quality of 6 or below are discarded.

Under the indel model, only reads that are considered informative for all possible indels of up to size 10bp (by default) are considered. For each event size up to the maximum to be considered, bases are removed from the reference to simulate a deletion or from the read to simulate an insertion. A read is considered informative if, for the pair of sequences with the indel modification, the sum of the base qualities for mismatching bases in the read is greater than the sum of the base qualities for bases that are mismatched according to the original alignment. The number of reads informative for indels is capped at 40. A constant reference quality of 45 is used for each indel-informative read.

Genotype likelihoods for genotypes incorporating the non-reference allele are calculated in the same way as described above for likelihoods derived from both SNP and indel models. The likelihoods with the lower genotype quality are then assigned to the site.

Output from the HC-RCM is captured in a new intermediate file format called genomic variant call format (gVCF) that contains likelihood data for every position in the genome (or specified intervals). This is an intermediate output that is not appropriate for analysis until the data has been processed with the full GATK Best Practices pipeline.

### 4.5 Running HaplotypeCaller-Reference Confidence Model

Use of the Reference Confidence Model is not the HaplotypeCaller default and must be specified on the commandline. (See https://software.broadinstitute.org/gatk/documentation/article?id=3893 for exact command lines.) Output from this mode will be intermediate gVCF files that are not appropriate for analysis. The reader is encouraged to visit the GATK Forums website for more information about the gVCF intermediate file format used by the HC-RCM (https://software.broadinstitute.org/gatk/documentation/article.php?id=4017). gVCFs must be geno-typed with the GATK GenotypeGVCFs tool to remove low quality, likely artifactual alleles and assign a quality score to each variant. For many of the scaling analyses presented here, per-sample gVCFs were first combined into multi-sample gVCFs using the GATK3 tool CombineGVCFs before genotyping that output with GenotypeGVCFs. For multi-sample analysis, Geno-typeGVCFs also produces the finalized annotation values that will be used as features in VQSR filtering.

